# Development and Validation of an LC-MS Method for Quantification of Sex Steroid Hormones in Skeletal Muscle

**DOI:** 10.64898/2026.05.12.724720

**Authors:** Viktor Engman, Séverine Lamon, Shaun Mason

## Abstract

Sex steroid hormones are not exclusively localised in the circulation and can be found in numerous extragonadal tissues, in concentrations unrelated to the circulating fraction. Existing methodology to measure intramuscular steroid hormone concentrations includes both immune-based assays and liquid chromatography-mass spectrometry (LC-MS), the gold standard for hormone measurements. To date, no LC-MS based methods validation has been published on the measurement of intramuscular sex steroid hormones, despite clear biological relevance. Here, we describe the development and validation of a simple, high-throughput LC-MS Orbitrap method for the measurement of 10 intramuscular sex steroid hormones, including pregnenolone, progesterone, dehydroepiandrosterone, androstenedione, testosterone, epitestosterone, dihydrotestosterone, oestrone, oestradiol, and oestriol. In brief, isotope labelled standards were added to 5-6 milligrams of lyophilised muscle tissue, homogenised and extracted with ethyl acetate. The extracts were dried down and sequentially derivatised with 1-methylimidazole-2-sulfonyl chloride and hydroxylamine hydrochloride to target both the phenolic hydroxyl groups and ketone groups. The limit of detection was 1.0 ± 1.0 pg/mg (range 0.36 - 3.26 pg/mg), with a R^2^ > 0.99 for all analytes. Matrix effects were 90-110% for all analytes except for dihydrotestosterone (143.6%), and precision was <10 CV% for all analytes in the presence of a muscle matrix. Our method allows for 20-40 samples to be prepared in ∼4 h, with a sample data acquisition time of 13 minutes. Moreover, our method provides the opportunity for specific analysis of steroid hormone concentrations in skeletal muscle, allowing target tissue specificity instead of relying on proxy measures from the circulation.

## 2 Introduction

Sex steroid hormones are potent endocrine signalling molecules synthesised from their common precursor cholesterol, primarily in the gonads and adrenal glands [1,2]. Upon secretion into the bloodstream, they are transported to their target tissues either tightly bound to binding proteins, loosely bound, or unbound, where the latter two constitute the bioavailable hormone fraction [3]. Unless tightly bound to binding proteins, steroid hormones can freely bind to their cytosolic receptors before translocating into the nucleus and modulate the expression of multiple genes [4,5]. One such target tissue is skeletal muscle, where, for example, the androgen receptor (AR) canonical signalling pathway enables testosterone to bind to the AR and upregulate genes governing muscle hypertrophy [6]. Similarly, oestrogens may enter the skeletal muscle cell, bind to the oestrogen receptor (ER), and modulate gene expression at the nucleus [7]. The progesterone receptor (PR) has also been detected in skeletal muscle tissue, but progesterone’s interaction with its receptor is less well understood [8], and it has been postulated that progesterone may be able to bind to the ER and antagonise the effects of the oestrogens [9]. Furthermore, pregnenolone, the first precursor in the sex steroid hormone synthesis pathway, lacks a specific nuclear receptor and its putative actions on skeletal muscle remains elusive [10].

Sex steroid hormones may also be synthesised in extragonadal tissue, including skeletal muscle [11–13]. The transcripts of 3β-hydroxysteroid dehydrogenase (*3β-HSD*), 17β-hydroxysteroid dehydrogenase (*17β-HSD*), aromatase (*P450arom*), 3-oxo-5α-steroid-4-dehydrogenase (*5α-reductase*) are all expressed in skeletal muscle, enabling local enzymatic conversion of steroid hormones [14,15]. Whilst translocator protein (*TSPO*), steroidogenic acute regulatory protein (*StAR*), and cytochrome P450 family 11 subfamily A member 1 (*CYP11a1*) have been detected in mouse and avian muscle [16], *StAR* and *CYP11a1* are not detectable in C2C12 cells and rat muscle [17–19], and are very lowly expressed in humans muscle [20]. This suggests that the ability for intramuscular *de novo* steroid hormone synthesis from cholesterol in skeletal muscle may not be conserved across species, and possibly indicates an increased importance of the cholesterol-independent sulfatase pathway for pregnenolone synthesis [17–19,21].

Previous research suggests that circulating and intramuscular sex hormones concentrations are not significantly associated and may be regulated independently [22]. This phenomenon is documented across various tissues including brain and adipose tissue [23,24], and may extend to skeletal muscle. Indeed, in a comparison of intramuscular oestradiol (E2) levels between pairs of postmenopausal twins in which one twin used oestrogen-containing hormone replacement therapy, intramuscular E2 concentrations was unaffected by treatment status [13]. Intramuscular hormone concentrations also appear to differ according to the fibre type composition of the muscle [16]. Intramuscular hormone status therefore cannot be inferred from blood-based measurements, altogether raising questions about its translatability to intramuscular hormonal status.

In clinical settings, a venous blood sample can be drawn and analysed to assess circulating hormone concentrations via well-established mass-spectrometry based methods [25]. Most previous research into the interplay between intramuscular sex hormones, ageing, and muscle mass and function has however been constrained by the use of immunoassays [26]. Whilst modern immunoassays demonstrate high accuracy and excel in simplicity, their measurement variability increases when tested at low hormonal concentrations compared to the gold-standard methodology mass spectrometry [27,28]. Notwithstanding the unknown effects of the muscle matrix, these assays also lack the ability to quantify multiple steroid hormones simultaneously.

Sex steroid hormones exert marked physiological effects even at low concentrations, which present further analytical challenges for accurate analysis. Whilst not required for liquid chromatography-mass spectrometry (LC-MS) analysis, chemical derivatisation with agents such as dansyl chloride and picolinic acid can enhance detection sensitivity in phenolic hydroxysteroids [29,30]. Similarly, derivatisation with 1-methylimidazole-2-sulfonyl chloride (1M2S) targeting oestrogens can increase assay sensitivity manyfold [29]. Likewise, ketosteroids have been successfully derivatised using hydroxylamine hydrochloride (HL), improving assay sensitivity [31]. Despite the cyclopentanoperhydrophenanthrene four-ring structure being largely identical in all sex steroid hormones [2], differences in functional groups pose a challenge for chemical derivatisation to increase the assay sensitivity for structurally different compounds. Therefore, there is a need to develop highly sensitive and integrated methods that can target a broad range of different steroid classes and their associated compounds within a single analytical run.

Whilst LC-MS has been used in a number of studies to measure intramuscular sex hormones [19,32–37], including in a methodological paper analysing hormone concentrations across multiple tissue**s** [38], there is a lack of analytical validation of LC-MS assays specifically targeting intramuscular sex steroid hormones. Despite the variable use of LC-MS, immunoassay-based methods remain commonplace for intramuscular sex hormone measurements [26]. Thus, we developed and validated a simple LC–MS based method using a two-step derivatisation protocol for the simultaneous quantification of ten sex steroid hormones in skeletal muscle tissue.

## 3 Methods

### 3.1 Ethics

Mouse skeletal muscle samples were obtained through our institutional tissue-sharing program from previously completed studies. All animal procedures were conducted in accordance with institutional and national guidelines and were approved by the Deakin University Animal Ethics committee. The use of shared tissues complied with the ARRIVE guidelines and institutional policies on secondary use of animal-derived materials [39]. As the present study involved the use of existing, de-identified tissues only, no additional animal ethics approval was sought.

Ethical approval was granted by the Deakin University Human Research Ethics committee (project number 2021-307) for the collection of the human muscle biopsies here used as representative samples. The biopsies were from a healthy male (32 years old) and healthy female (38 years old). Ethical approval was granted by the Deakin University Animal Ethics committee (project number 2024/AE000058) for the collection of mouse tissues here used as representative samples. These were from a sedentary male (16 weeks old) and a sedentary female (16 weeks old) C57BL/6 mouse.

### 3.2 Materials

Neat isotope labelled steroid hormones standards dihydrotestosterone (DHT-d_4_; Novachem, Cat. No. DLM-9041-0.001), epitestosterone (EpiT-d_3_; Novachem, Cat. No. D548), oestrone (E1-d_2_; Sapphire Bioscience, Cat. No. 9002844), oestriol (E3-d_2_; Sapphire Bioscience, Cat. No. 9002844), progesterone (P4-d_9_; Sapphire Bioscience, Cat. No. 25047) were prepared to 1 mg/mL in methanol, whereas the remaining stock solution came pre-suspended: androstenedione (A4-^13^C_3_; 100 μg/mL in acetonitrile, Novachem, Cat. No. A-084-1ML), dehydroepiandrosterone (DHEA-d_5_; 100 μg/mL in methanol, Novachem, Cat. No. D-064-1ML), oestradiol (E2-d_5_; 100 μg/mL in acetonitrile, Novachem, Cat. No. E-061-1ML), pregnenolone (P5-^13^C_2_, d_2_; 100 μg/mL in acetonitrile, Novachem, Cat. No. CDLM-9158-C), and testosterone (T-d_3_; 100 μg/mL in 1,2-dimethoxyethane, Merck, Cat. No. T5536-1ML), and all stored at -20°C. Neat unlabelled steroid hormone standards oestrone (E1; Merck, Cat. No. PHR1535-500MG), oestradiol (E2; Merck, Cat. No. PHR1353-1G), oestriol (E3; Merck, Cat. No. E-074), progesterone (P4, Sapphire Bioscience, Cat. No. 15876), and testosterone (T; Merck, Cat. No. 46923-250MG-R) were prepared to 1 mg/mL in methanol whereas the remaining stock solution came pre-suspended: androstenedione (A4; 1 mg/mL in acetonitrile, Novachem, Cat. No. A-075-1ML), dehydroepiandrosterone (DHEA; 100 μg/mL in methanol, PM Separations, Cat. No. C9000.19-100-ME), dihydrotestosterone (DHT; 1 mg/mL in methanol, Merck, Cat. No. D-073), epitestosterone (EpiT; 1 mg/mL in acetonitrile, PM Separations, Cat. No. C13179.19-K-AN), pregnenolone (P5; 100 μg/mL, Novachem, Cat. No. P-104-1ML), and all stored at -20°C.

All standards were diluted to 1 µg/mL in methanol. These were then used to prepare the standard cocktails. The internal standard cocktail was mixed to 10 pg/μL for each analyte, except for the oestrogens which were mixed to 5 pg/ μL, and 10 μL was added to each sample. Nine unlabelled standard cocktails were prepared to 0, 0.1, 1.0, 2.5, 5, 10, 25, 50, and 100 pg/μL per steroid, respectively, and 10 μL was added to the corresponding sample. All solvents used in standard and sample preparation were LC-MS grade purity.

### 3.3 Sample preparation

Standard curves were generated both in the presence and absence of a muscle matrix. Approximately 1 gram of gastrocnemius mouse muscle was lyophilised in a 15 mL falcon tube for ∼16 hours overnight in a freeze drier. Immediately upon removal from the freezer, five stainless-steel beads (5mm; Qiagen Cat. No. 69989) were added and the tube was vigorously shaken until the muscle tissue had formed a homogenous powder. The muscle powder was then aliquoted into nine 2 mL screw-top tubes (∼5.2 mg for each tube) forming the muscle matrix for the nine standards, and a stainless-steel bead (5mm; Qiagen, Cat. No. 69989) was added. Ten microlitres of the internal standard cocktail, 10 μL of the corresponding unlabelled standard cocktail, and 1 mL ethyl acetate pre-cooled in ice were combined with the samples and then homogenised for 2 x 1 min on a precooled (at -20°C) 24 tube block on a TissueLyser III (Qiagen, Cat. No. 9003240). Thereafter, 600 μL of sodium acetate (200mM, pH 5; pre-cooled in ice) was added and the samples, which were placed in a ThermoMixer for 10 mins (4°C, 2,000 RPM) and then centrifuged for 10 mins (4°C, 10,000g). Then, 800 μL from the top layer was extracted and transferred to a new 2 mL tube. Another 800 μL of ethyl acetate was then added to the original tube, which was placed in the ThermoMixer for 2 mins (4°C, 2,000 RPM) and then centrifuged for 10 mins (4°C, 10,000g). Another 800 μL was then taken from the top layer and added to the first 800 µL extract. The total 1,600 μL of extract was then dried using a CentriVap vacuum concentrator (Labconco) set to 40°C for ∼40 minutes (until completely dry).

### 3.4 Derivatisation

The derivatisation was firstly carried out by adding 50 μL sodium bicarbonate (NaHCO_3_; 100mM, pH 9, Merck, Cat. No. 13433) and 50 μL of 3 mg/mL 1-methylimidazole-2-sulfonyl chloride (1M2S; Merck, Cat. No. 718211) in acetone and incubated at 60°C for 15 minutes. The samples were then dried using the CentriVap vacuum concentrator (Labconco) set to 40°C for ∼70 mins until completely dry. The residue then underwent a second derivatisation step with 7 mg/mL hydroxylamine hydrochloride (HL; Merck, Cat. No. 159417) in 30% acetonitrile for 30 minutes of incubation at 60°C. The samples where then transferred to autosampler vial inserts and stored at 10°C in the autosampler. Twenty-five microlitres was then injected onto the LC-MS.

### 3.5 Liquid chromatography and mass spectrometry

The analysis was carried out on a Vanquish Flex LC coupled to an Orbitrap Exploris 240 (Thermo Fisher) using an Ascentis® Express 90 Å C18 (2.7 μm) HPLC Column guard column (Merck, Cat. No. 53501-U) in a Ascentis® Express Guard Cartridge Holder (Merck, Cat. No. 53500-U) coupled to an Ascentis® Express 90 Å C18, 2.7 μm HPLC column (Merck, Cat. No. 53825-U). The mobile phases consisted of 0.1% formic acid in water (A) and 0.1% formic acid in acetonitrile (B) with a flow rate of 500 μL/min and a linear gradient (A/B) of 65/35–47.5/52.5 from 0 to 7.5 mins, then 0/100 from 7.5 to

9.5 mins and then finally to 65/35 for 3.5 mins. A switcher valve diverted the flow away from the Orbitrap outside the range of 1.0 - 7.4 mins. Mass detection was undertaken in positive electrospray ionisation mode with ion spray voltage set to: 4500V, sheath gas: 30, auxiliary gas: 5, sweep gas: 1, ion transfer tube temperature: 350°C, and vaporizer temperature: 320°C. Data acquisition was conducted in both full-scan mode and targeted selected ion monitoring (tSIM) using Xcalibur software (ThermoFisher Scientific). For the full scan, the range was set to m/z 280-440, RF lens 70%, and an orbitrap resolution of 30,000 Hz. For the tSIM, the isolation window was set to 2 m/z, RF lens 70%, and an orbitrap resolution of 45,000 Hz. All samples were run in triplicate.

### 3.6 Data analysis

Data analysis was conducted using Skyline (MacCoss Lab Software) where the tSIM trace was used for quantification with the ion match tolerance set to 5ppm [40]. The peak areas were then integrated into Microsoft Excel where peak area was calculated as target peak area divided by corresponding internal standard peak area, multiplied by the internal standard added (in picograms), and then divided by the lyophilised muscle mass (in milligrams) in the sample [41].

Limit of detection (LOD) was calculated as 3.3*(*SD_intercept_* / *x̅_slope_*) and limit of quantitation (LOQ) was calculated as 10*(*SD_intercept_* / *x̅_slope_*) [42]. Precision was calculated at three levels (low, mid, high) and presented as the coefficient of variation (*SD*/*x̅*) [42]. In the absence of intramuscular hormone data, we defined the *low* condition as the lowest unlabelled standard concentration that resulted in confirmed peaks in all three technical replicates and ≥LOD. The *mid* and *high* condition represented the next two higher standard concentrations with confirmed peaks in all technical replicates. The matrix effect was calculated as the slope of the standard curve with a muscle matrix divided by the slope of the standard curve in the absence of a muscle matrix across the linear range [42].

## 4 Results and Discussion

We developed and validated a two-step derivatisation strategy to quantify intramuscular sex hormone concentrations of androgens A4, DHEA, DHT, EpiT and T; oestrogens E1, E2, and E3; and progestogens P4 and P5. During method development, we investigated using HL as a single derivatisation agent. However, we were unable to detect E2 and E3 with sufficient sensitivity using HL alone. With the novelty of intramuscular sex hormone research, we wanted to include E2, the main female hormone, as well as E3, despite typically associated with pregnancy [1], to enable exploratory work. Likewise, when we only used the derivatisation agent 1M2S, oestrogens were detected with good sensitivity, but androgens and progestogens were not. By sequentially derivatising the extracts with 1M2S followed by HL, all oestrogens were first derivatised with 1M2S, and all androgens and progestogens were then derivatised with HL. E1 was the only analyte that formed a double derivatised product, where 1M2S interacts with the position 3 hydroxyl group and HL interacts with the position 17 ketone group. The target analytes and their corresponding stable isotope labelled standards, retention times, and interactions with derivatisation agents have been outlined in Table 1.

**Table 1.**
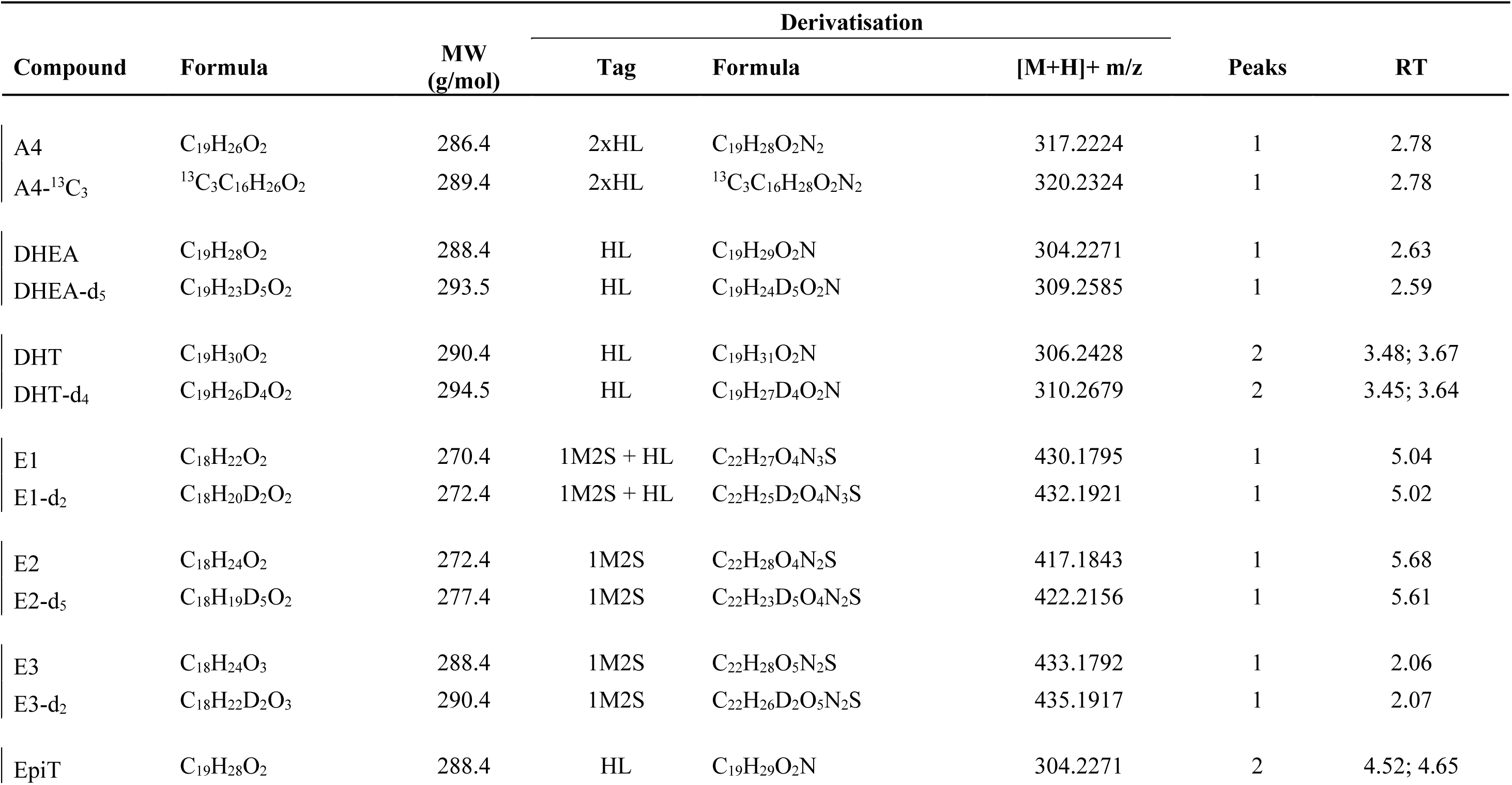
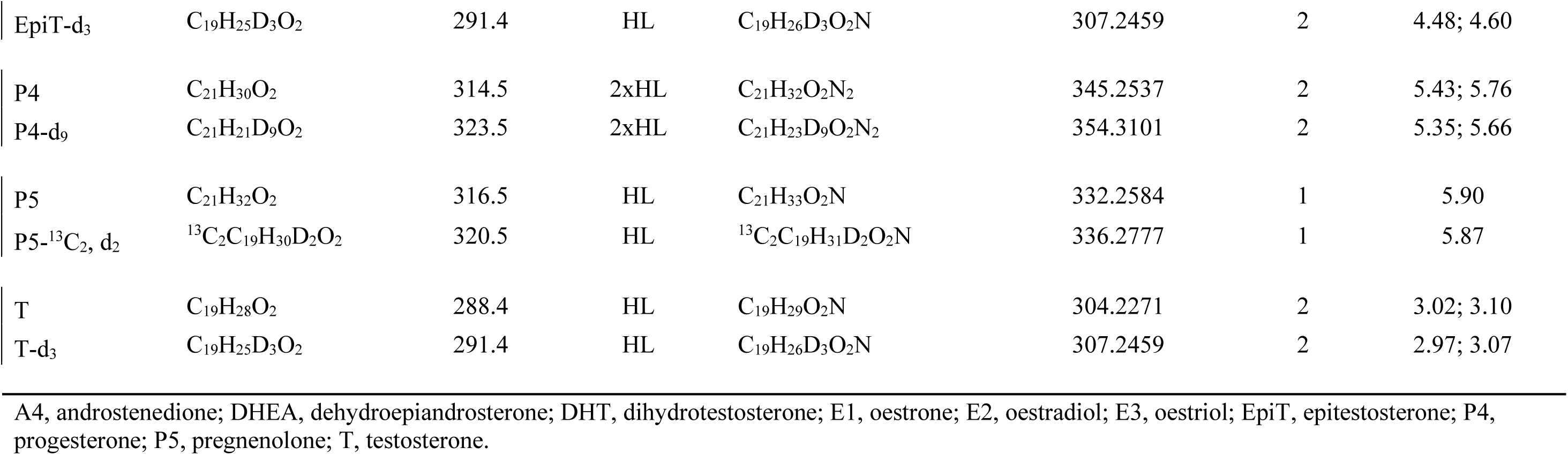
Compound details.

With our two-step derivatisation method, sample preparation took ∼4 hours for 20-40 samples, primarily limited by equipment capacity (e.g., centrifuge batch size), followed by 13 mins of LC-MS acquisition time per sample. The run plan is outlined in Figure 1, where all analytes eluted between 2 – 6 minutes. Whilst 1M2S yielded singlet peaks for all target analytes, HL resulted in the formation of stereoisomers for DHT, EpiT, P4, and testosterone, consistent with previous findings [31,43]. Both peaks were integrated for area for testosterone and DHT, whereas the second larger peak were used for EpiT and P4. Whilst the stereoisomers were well separated for P4, there was some coelution of the two EpiT peaks. Nevertheless, single peak integration allowed for clear peak identification at lower concentrations than double peak integration for EpiT. Synchronised integration was used for DHEA, which included peak integration of the same time window relative to the peak retention time for both the unlabelled and labelled standards. For the remaining peaks, the Skyline default automatic integration was used and adjusted manually where appropriate upon subsequent visual inspection.

**Figure 1.**
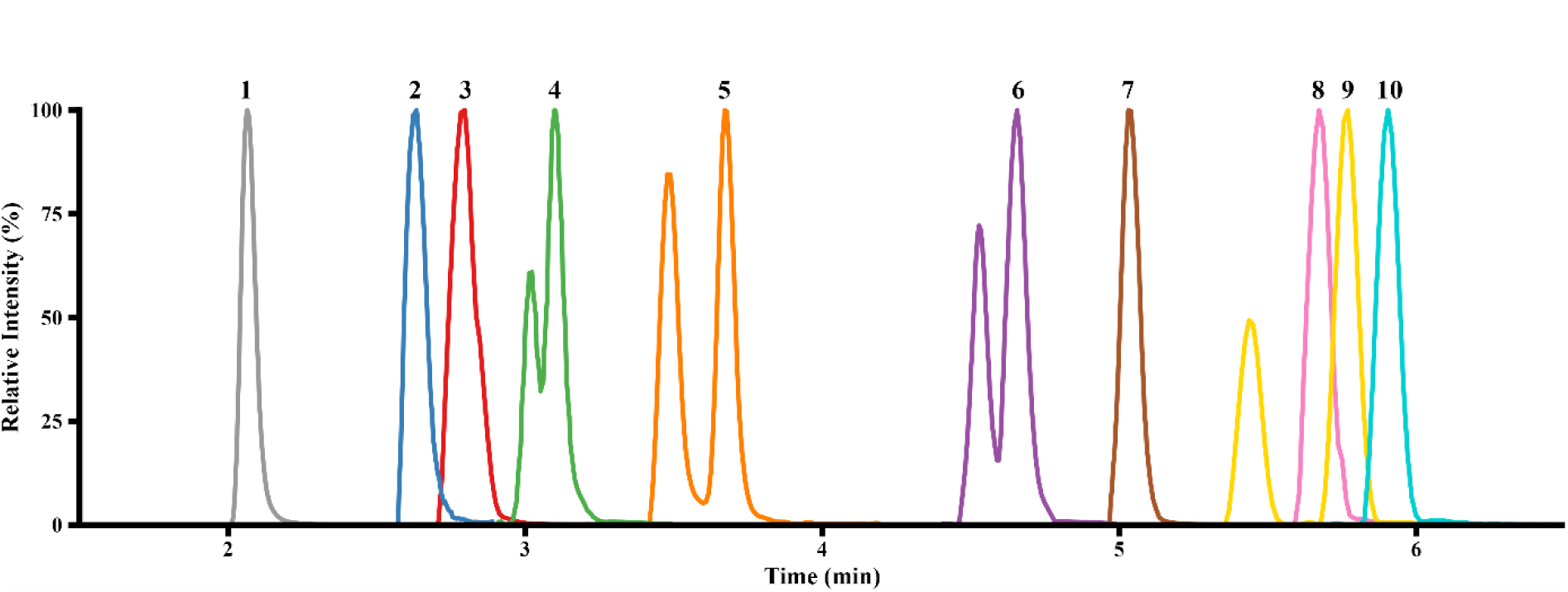
LC-MS run plan. (1) is E3, (2) is DHEA, (3) is A4, (4) is testosterone, (5) is DHT, (6) is EpiT, (7) is E1, (8) is E2, (9) is P4, (10) is P5.

Sample carryover was assessed by injecting a blank sample after the injection of supraphysiological levels of the unlabelled standards (1,000 pg unlabelled standards added) in the presence of a muscle matrix, and no carryover was detected. To assess the linear range, calibration standards were added in both the presence and absence of a muscle matrix, with the full results outlined in Figure 2 and Figure 3. R^2^ > 0.99 was achieved for all analytes in both conditions. The main limitation for the validation process was the absence of a hormone free muscle matrix. Using muscle from ovariectomised mice does not remove the intramuscular hormone fraction [44], hormone free muscle tissue is not commercially available, and repurposing strategies from blood-based research, such as charcoal stripping, would have prevented the addition of standards to a solid matrix. As the muscle was not devoid of sex steroid hormones, a standard curve in the presence of the muscle matrix could only be constructed at levels exceeding the already present endogenous hormones (Figure 2).

**Figure 2.**
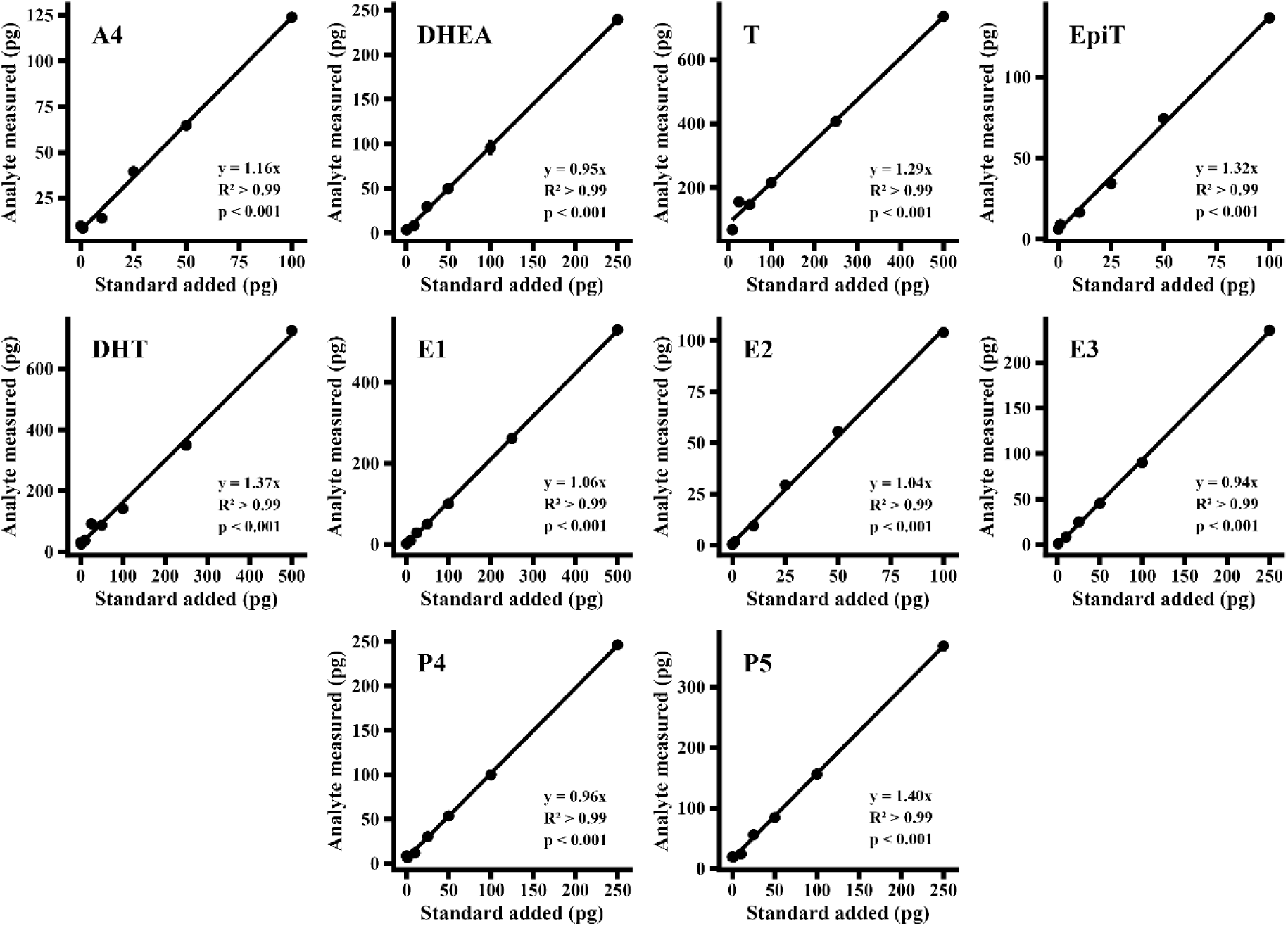
Muscle standard curves. All analytes had a R^2^ > 0.99 and p < 0.001.

**Figure 3.**
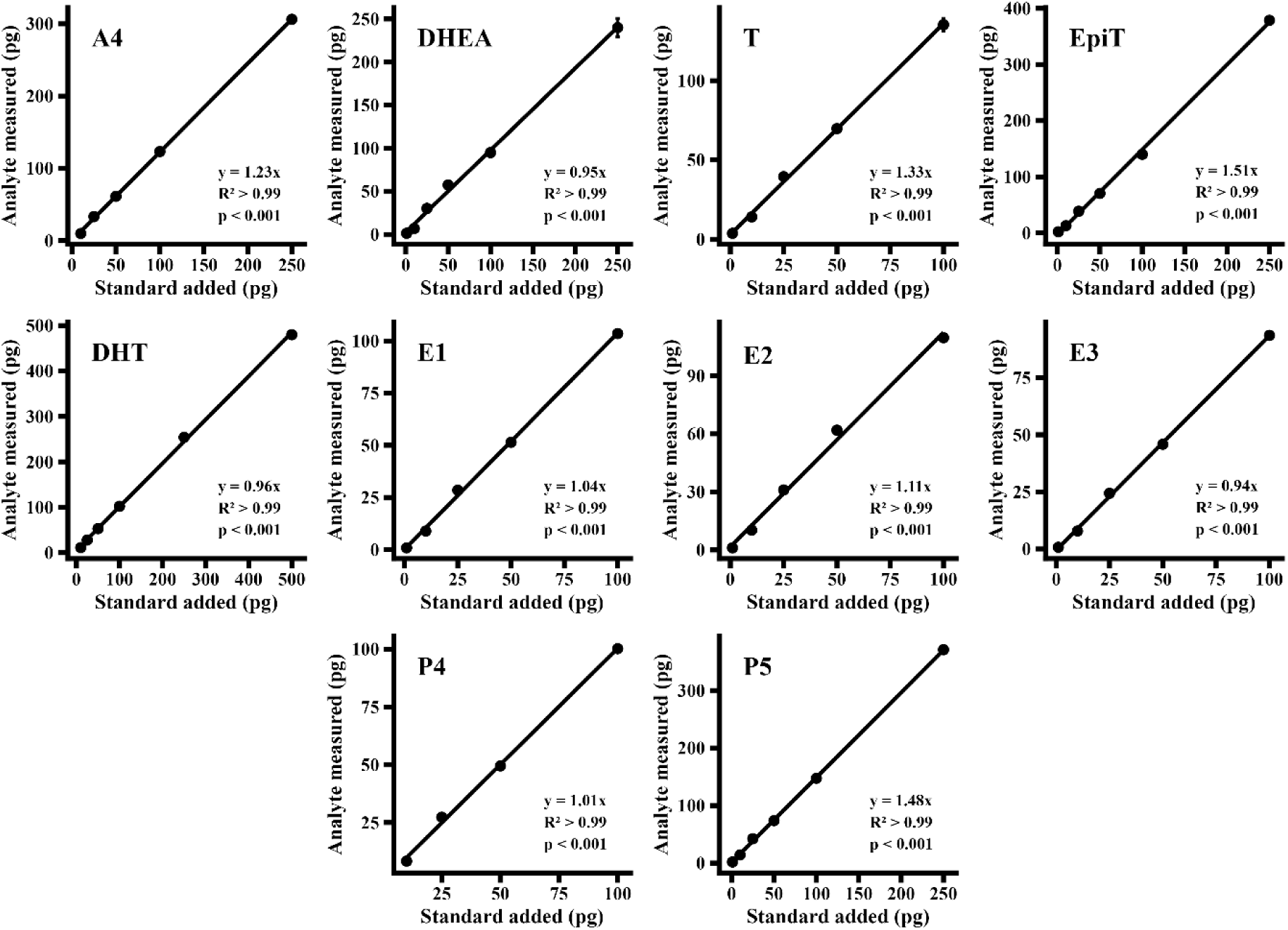
Solvent standard curves. All analytes had a R^2^ >0.99 and p < 0.001.

LODs and LOQs were calculated as 3.3 and 10.0 multiplied by the standard deviation for the y-intercept divided by the average slope [42]. LOD was 1.0 ± 1.0 pg/mg (range 0.36 - 3.26 pg/mg) and LOQ was 3.0 ± 2.9 pg/mg (range 1.10 – 9.89 pg/mg) when extracted from ∼5.2 mg of lyophilised muscle tissue (full results in Table 2 and Table 3). For the analytes with detectable hormone levels in the muscle matrix, this would have elevated the y-axis intercept and restricted the linear range, most notably for testosterone which returned the highest LOD (3.26 pg/mg) and LOQ (9.89 pg/mg) with a linear range beginning at 13.2 pg/mg. At low spike concentrations, the quantification of the unlabelled standards may be partially obscured by the endogenous background, potentially reducing the reliability of those calibration points. Nevertheless, linearity persisted still in the sub-physiological ranges when unlabelled standards were added to a solvent in the absence of a muscle matrix (Figure 3).

**Table 2.**
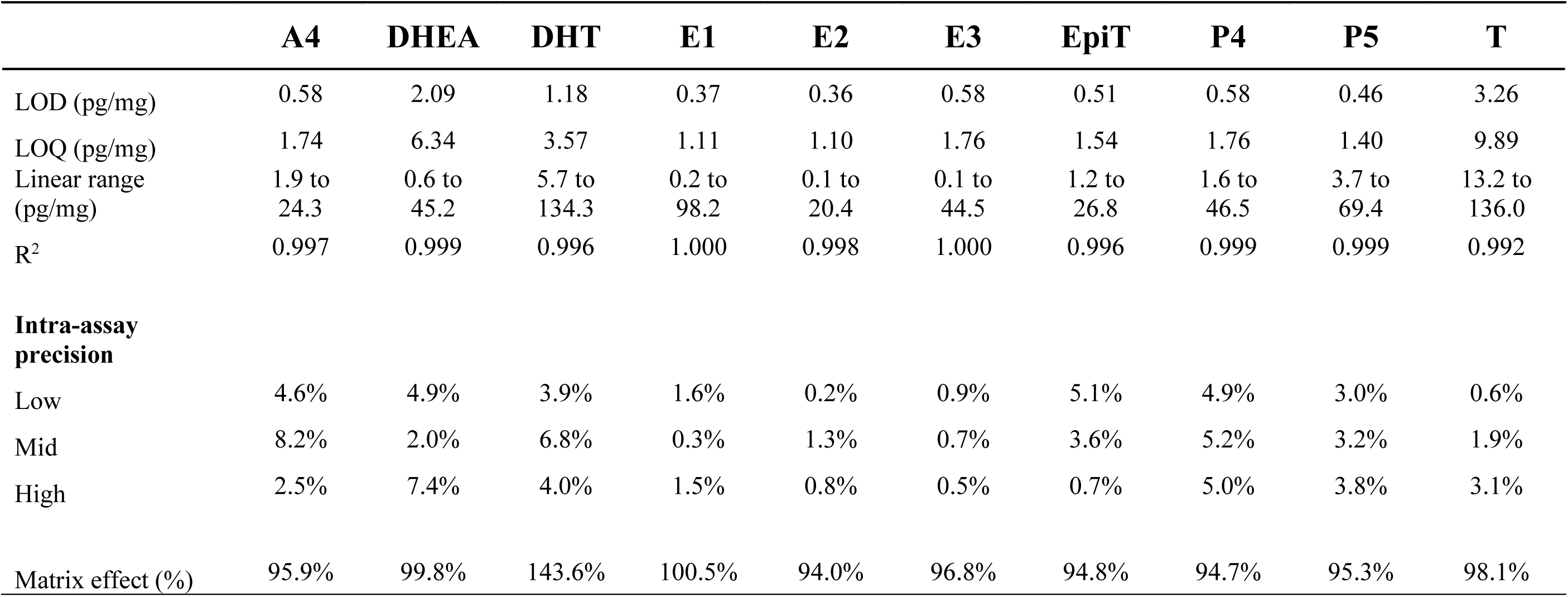
Methods performance in muscle. LOD, LOQ, and linear range are given as picograms (pg) of analyte per milligram (mg) of lyophilised muscle tissue. Intra-assay precision refers to the coefficient of variation (CV%) of three technical replicates at the lowest sample with a concentration equal to or higher than LOD (Low), and the two upper levels were the next two vials with higher concentrations.

**Table 3.** Methods performance in solvent. LOD, LOQ, and linear range are given as picograms (pg) of analyte. Intra-assay precision refers to the coefficient of variation (CV%) of three technical replicates at the lowest sample with a concentration equal to or higher than LOD (Low), and the two upper levels were the next two vials with higher concentrations.

|  | A4 | DHEA | DHT | E1 | E2 | E3 | EpiT | P4 | P5 | T |
| --- | --- | --- | --- | --- | --- | --- | --- | --- | --- | --- |
| LOD (pg) | 1.14 | 5.06 | 1.14 | 0.99 | 1.00 | 0.86 | 0.66 | 1.26 | 0.33 | 2.35 |
| LOQ (pg) | 3.45 | 15.34 | 3.45 | 3.00 | 3.02 | 2.61 | 1.99 | 3.82 | 0.99 | 7.13 |
| Linear range (pg) | 10.0 to 250.0 | 1.0 to 250.0 | 10.0 to 500.0 | 1.0 to 100.0 | 1.0 to 100.0 | 1.0 to 100.0 | 1.0 to 250.0 | 10.0 to 100.0 | 1.0 to 250.0 | 1.0 to 100.0 |
| R<sup>2</sup> | 1.000 | 0.997 | 0.999 | 0.998 | 0.994 | 0.999 | 0.999 | 0.998 | 1.000 | 0.998 |
| **Intra-assay precision** |  |  |  |  |  |  |  |  |  |  |
| Low | 1.4% | 7.5% | 5.8% | 0.6% | 1.5% | 1.1% | 9.4% | 5.1% | 14.7% | 9.7% |
| Mid | 1.3% | 1.7% | 3.9% | 0.4% | 0.5% | 0.9% | 5.8% | 3.3% | 7.5% | 4.4% |
| High | 1.7% | 6.5% | 1.6% | 3.6% | 2.7% | 1.1% | 0.5% | 0.5% | 3.4% | 2.6% |

Optimised LC-MS methods for routine use in plasma commonly report lower LODs and LOQs than our method [27], which is possibly more attributable to matrix homogeneity and larger extraction volumes than analytical sensitivity, rendering direct cross matrix comparisons of limited utility. Nouri *et al.* demonstrated that extracting hormones from 250 μL compared to 10 μL plasma resulted in a 20-fold lower LOD for testosterone, 40-fold lower LOD for P5, and 2-fold lower LOD for E1 [45]. Nevertheless, LOD and LOQ shall not be interpreted as definitive limits as these may vary depending on how much tissue was used for extraction, as well the estimation method used [46].

Precision was deemed high, with all CVs <10%, except for P5 in solvent in the *low* condition, which was 14.7%. Overall, the analytes derivatised with 1M2S had a higher precision (all < 4% in both muscle and solvent) than the analytes derivatised by HL. Matrix effect was between 90-110% for all analytes except for DHT, where it was 143.6% (Table 2 & Table 3), which is similar to previously published methodological studies on LC-MS based hormone measurements [47,48].

Heterogeneity in quantification methodology and what the hormone levels have been normalised to for muscle tissue in previous research limits comparability of results between studies. As we opted to normalise hormone levels to lyophilised muscle tissue instead of protein content or non-lyophilised mass, comparisons to previous research were not possible. Harmonisation is therefore warranted, where we suggest normalising hormone levels to lyophilised muscle, to minimise the influence of variability in tissue water content. Sample hormone data from a male and female, as well as from the soleus, extensor digitorum longus, and diaphragm muscle from a male and female mouse have been outlined in Table 4.

**Table 4.**
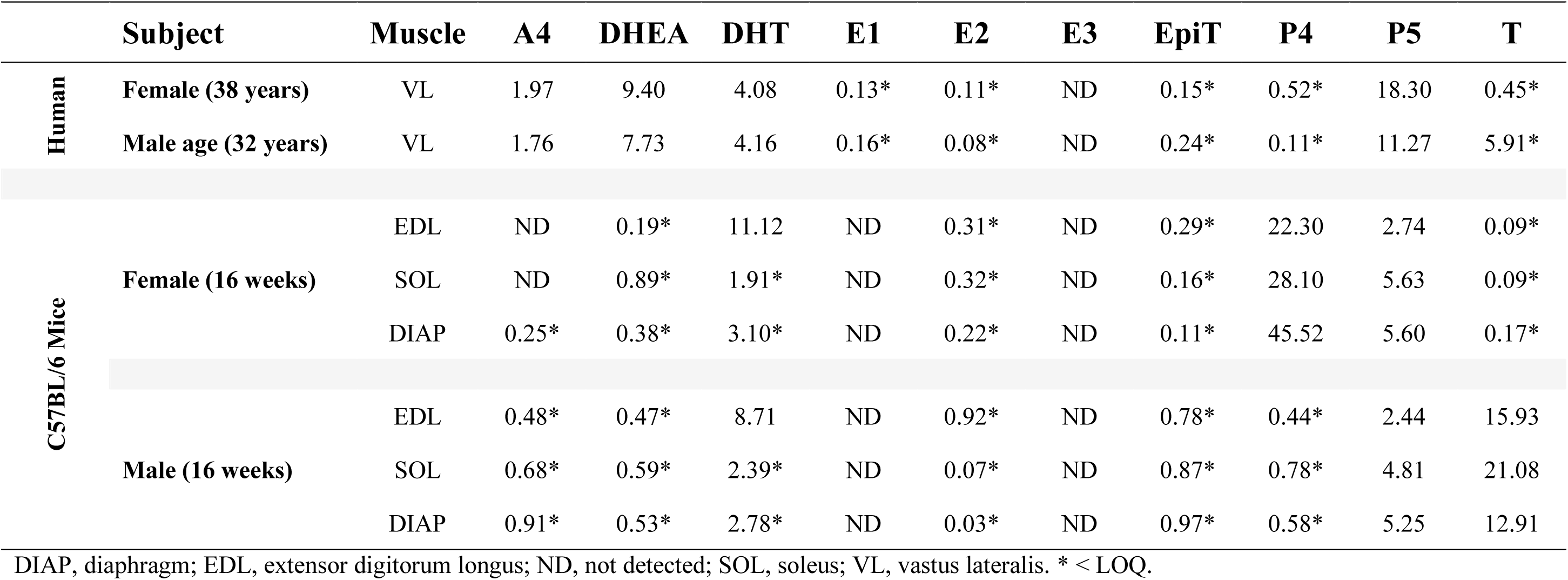
Intramuscular hormone data from human and mouse muscle. Hormone concentrations are expressed as picograms (pg) of analyte per milligram (mg) of lyophilised muscle tissue.

Whilst our method provides insight into the present state of intramuscular sex hormone concentrations, it cannot discriminate between the origin of these molecules. Similar to other tissues, skeletal muscle expresses the required enzymes to synthesise sex steroid hormones from their precursors [11,16]. As the intramuscular hormone fraction appears uncorrelated to the corresponding circulating hormone fraction [22], it is reasonable to assume that at least some fraction of the intramuscular hormones measured were synthesised in the muscle tissue.

Although residual blood contamination cannot be completely excluded, several lines of evidence indicate that it does not meaningfully contribute to the intramuscular hormone signal. First, within the same extensor digitorum longus samples from sedentary male C57BL/6 mice, intramuscular–plasma relationships were hormone-specific, ranging from very strong (testosterone: ρ = 0.930, *p* < 0.001, *n* = 12) to absent (DHT: ρ = 0.035, *p* = 0.921, *n* = 12; data not shown). Second, a simple mass-balance consideration demonstrates that the intramuscular DHT concentrations observed cannot be explained by plasma carryover: doing so would require implausible plasma fractions exceeding 100% of tissue volume in the majority of samples (∼75%). These findings collectively indicate that the measured intramuscular hormone concentrations predominantly reflect local tissue content rather than residual blood.

Validation was performed using an Orbitrap Exploris 240. Whilst Orbitrap instruments are well-suited for exploratory work, triple quadrupole platforms generally outperform Orbitraps for targeted quantification of low-abundance compounds [49]. Therefore, validation with a triple quadrupole-based method is warranted for routine work of intramuscular hormone concentrations, specifically in samples with low endogenous concentrations. However, such validation would remain constrained by the lack of a hormone free muscle matrix. Some tissue-based studies have used NaCl solutions as a surrogate hormone free matrix [43,50]. However, saline does not mimic the biological complexity of muscle tissue, which includes proteins, lipids, and other endogenous compounds all affecting ionisation. Therefore, the development of a hormone free muscle matrix is a prerequisite for robust tissue-specific methods validation.

## 5 Conclusion

Whilst the heterogenous and replete nature of skeletal muscle poses challenges for accurate method validation, we developed and validated a time-efficient and high-throughput LC-MS based method for sensitive simultaneous quantification of androgens, oestrogens, and progestogens, all within the same 13-minute analytical run from skeletal muscle tissue. This approach overcomes the limitations of blood-borne proxy measures, instead providing direct insight into the intramuscular sex hormone concentrations.

## 6 CRediT Statement

V.E. was involved in the conceptualisation, data acquisition, data analysis, and writing and revising the manuscript. S.L. was involved in conceptualisation, revising the manuscript, and funded the study. S.M. was involved in conceptualisation, data analysis, revising the manuscript, and funded the study.

